# Human Gut Phageome Analysis Uncovers Thousands of Highly Modular Endolysins

**DOI:** 10.64898/2026.04.25.720813

**Authors:** Raphael Kabir Niloy, Nurnabi Azad Jewel, Daniyal Karim, Mohimenul Haque Rolin, Tahsin Khan, Arzuba Akter, Shakhinur Islam Mondal

## Abstract

The escalating threat of antimicrobial resistance has renewed global interest in bacteriophages as precise and powerful tools for controlling bacterial populations in the human gut. These viruses owe much of their antibacterial potential to phage-encoded endolysins, enzymes capable of rapidly degrading bacterial cell walls with high specificity and low potential for resistance development. Despite their therapeutic promise, the overall composition of the gut phageome and the structural modularity of its endolysins remain poorly understood. In this study, we performed a large-scale analysis of 9,141 human gut metagenomic samples from 34 independent studies. Using standardized workflows for assembly, genome clustering, host prediction, and protein domain annotation, we reconstructed 15,267 phage genomes and identified 3,794 corresponding endolysins. The recovered genomes showed substantial variation in size and coding density, with an average GC content of 43%. Host prediction indicated that most phages targeted bacterial members of the phyla *Bacillota* (41%) and *Bacteroidota* (23%). Endolysin sequences grouped into 296 protein families and displayed striking domain modularity. Catalytic domains such as Amidase_2 and Glyco_hydro_25 frequently co-occurred with cell wall–binding motifs including LysM and CW_7. Remarkably, one endolysin contained 15 distinct domains, the highest natural domain diversity reported to date. Collectively, this study represents the most comprehensive characterization of the human gut phageome and its encoded endolysins to date. The exceptional modular diversity uncovered highlights the gut phageome as a rich reservoir of endolysin variants, providing a strong foundation for developing next-generation therapeutics against multidrug-resistant bacterial pathogens.

## 1. Introduction

Antimicrobial resistance (AMR) has become one of the most critical global health threats, responsible for an estimated 1.27 million deaths annually and contributing to nearly 5 million additional deaths worldwide (UN, 2023). The World Health Organization predicts that, by 2050, AMR could claim up to 10 million lives each year (C. J. Murray et al., 2022). Beyond its direct health impacts, AMR undermines global efforts toward the Sustainable Development Goals (SDGs), jeopardizing food security, increasing poverty, and imposing an enormous economic burden—particularly in developing nations, where gross domestic product (GDP) losses may reach 5–7% annually (Anderson et al., 2019; World Bank, 2017). These alarming statistics highlight the urgent need for effective alternatives to conventional antibiotics to combat resistant bacterial infections.

Among emerging alternatives, phage-based therapy has gained renewed interest as a targeted and eco-friendly approach to controlling bacterial pathogens. Bacteriophages, the most abundant biological entities on Earth, specifically infect bacteria and have been used as therapeutic agents long before the antibiotic era (Wittebole et al., 2014). However, recent research has shifted focus toward phage-derived enzymes, particularly endolysins, which hold exceptional promise as next-generation antimicrobials. Earlier studies have demonstrated that endolysins could lyse most bacterial cells, including those that are resistant to antibiotics (Abdelkader et al., 2019; Harhala et al., 2018; Loeffler et al., 2003; Premetis et al., 2023).

Endolysins are peptidoglycan hydrolases that degrade bacterial cell walls, leading to rapid and specific bacterial lysis (E. Murray et al., 2021). The structure of endolysins is modular, consisting primarily of two components: the enzymatically active domain (EAD) and the cell wall-binding domain (CBD). EADs function as a catalyst to cleave specific bonds in the bacterial cell wall. On the other hand, the CBDs target the constituents of cell walls for the localization of the EADs. The domains are linked by a flexible connector, enabling the enzyme to effectively identify and decompose the bacterial cell wall (Mondal et al., 2020). Compared to traditional antibiotics, they offer several key advantages: (1) high specificity, reducing damage to commensal microbiota; (2) rapid bactericidal action independent of bacterial growth phase; (3) low likelihood of resistance development; (4) modular architecture that enables protein engineering for improved activity and host range; (5) demonstrated synergistic effects when combined with conventional antibiotics, and (6) non-toxic behavior to animal bodies. (Fujimoto et al., 2020; Pottie et al., 2024; Rahman et al., 2021; Rodríguez-Rubio et al., 2013; Walsh et al., 2021). These features make endolysins particularly attractive for combating multidrug-resistant pathogens such as *Staphylococcus aureus*, *Acinetobacter baumannii*, and *Pseudomonas aeruginosa* (Haddad Kashani et al., 2018).

The human gut microbiome represents one of the richest natural reservoirs of bacteriophages and their encoded endolysins (Georgakis et al., 2025). Within this complex ecosystem, bacteriophages play pivotal roles in maintaining microbial balance and influencing host health. The human gut virome is remarkably diverse, with phages continuously co-evolving with their bacterial hosts (Ofir & Sorek, 2018). Consequently, it serves as an untapped resource for discovering novel endolysins with unique domain architectures and functional diversity (Castro-Mejía et al., 2015). Understanding this intricate network of phage–bacteria interactions not only expands our knowledge of gut microbial ecology but also provides a foundation for developing phage-derived therapeutics against antibiotic-resistant infections.

Advances in metagenomic sequencing have revolutionized our ability to explore the hidden endolysin diversity (Georgakis et al., 2025). Through large-scale metagenomic analyses, researchers can reconstruct phage genomes, predict host associations, and identify genes encoding antimicrobial enzymes such as endolysins—without relying on culture-based methods (Fernández-Ruiz et al., 2018; Handelsman, 2004; Paul et al., 2025). Alongside conventional sequence-based search methods, novel algorithms and methodologies, including DeepMineLys (Fu et al., 2024), EnzymeMiner (Hon et al., 2020), and FoldSeek (Heinzinger et al., 2024), provide swift and more precise structure-oriented investigations in metagenomic databases. Such approaches enable the systematic characterization of the endolysins from the gut phageome. Notably, analyzing the domain composition and modularity of endolysins derived from human gut phages can provide crucial insights into their evolutionary adaptation and therapeutic applicability (Pottie et al., 2024). Although there are databases of enzybiotics and phage lysins like Enzybase (Wu et al., 2012), phiBIOTICS (Hojckova et al., 2013), GMEnzy (Wu et al., 2014), and PhaLP (Criel et al., 2021), there is a need for more comprehensive and specialized repositories (Bałdysz et al., 2024). Thus, despite the immense potential of gut phage endolysins, comprehensive databases cataloging their diversity, functional properties, and evolutionary relationships remain limited. This knowledge gap hinders the rational selection and engineering of endolysins for therapeutic development.

The creation of systematic, well-annotated endolysin databases is therefore essential for advancing both fundamental research and translational applications. Such resources would facilitate: (1) rapid identification of candidate endolysins with desired biochemical properties (Plotka et al., 2015); (2) comparative genomic analyses to understand evolutionary relationships and functional constraints (Oliveira et al., 2013); (3) structural and functional predictions to guide protein engineering efforts (Krishnappa et al., 2023); (4) discovery of novel endolysins with improved specificity or activity against emerging resistant pathogens (Fu et al., 2024); and (5) integration with clinical and epidemiological data to prioritize candidates for therapeutic development (Yoda et al., 2024). Furthermore, publicly accessible databases promote scientific transparency, enable collaborative research, and accelerate the transition from discovery to clinical application.

In this study, we performed a comprehensive metagenomic analysis of the human gut phageome to identify and characterize phage-encoded endolysins. We analyzed 9,141 metagenomic samples from 34 independent human gut datasets to reconstruct high-quality phage genomes, predict their bacterial hosts, and annotate endolysin domains. This study presents the first extensive catalog of endolysins derived from the human gut virome, uncovering their remarkable modular diversity and emphasizing the gut microbiome as a natural reservoir of bioactive enzymes with potential therapeutic applications against antimicrobial-resistant bacteria.

## 2. Methods

### 2.1 Retrieving Assembled Human Gut Metagenomic Data

Gut metagenomic data were retrieved from MGnify (Gurbich et al., 2023). The data was filtered for Host-associated biome > Human > Digestive System > Large Intestine. All the assembled sequences were screened with the Assembly (Experiment type) option. The metadata of the obtained studies were retrieved through the MGnify API with a Python script. After that, another Python script was used to select the analyses with the description “Processed Contigs” and retrieved them through the MGnify API. All these contigs were assembled by the EBI-Metagenomics (EMG) pipeline. Any sample was excluded if it was not human gut metagenomic data.

### 2.2 Preprocessing of Contigs

All the contigs under 1 kb in length were filtered out. After that, databases were created with the remaining sequences. The remaining sequences were searched against these databases for circular viral contigs prediction. If there was a 100% identity between the first and last 50 bases in a contig in the 5′ and 3′ end with at least 1.5 kb of contig length, then it was considered for further analysis for a circular viral contig (Fujimoto et al., 2020). This step was done using megablast flag in BLAST+ v.2.15.1 (Morgulis et al., 2008). For linear viral contigs with a length ≥ 5 kb were extracted. For viral contig prediction, all the pre-processed contigs were predicted using VirFinder (v1.1) with default settings (Ren et al., 2017). Only the sequences with p < 0.05 were retained.

### 2.3 Genome Completeness and Bacterial Decontamination

Viral genome completeness was assessed with CheckV using end-to-end mode (Nayfach et al., 2020). The CheckV database v1.5 was used. Sequences with no viral genes, <5% contamination, and low-quality genome fragments (<50% completeness) were excluded (Parks et al., 2015). Additionally, sequences for which quality could not be determined and proviral sequences were taken for the next step. CD-HIT-EST (Li & Godzik, 2006) was used to remove redundancy, where 95% similar sequences were filtered out (-c 0.95 -G 1 -n 10 - mask NX). All the sequences passing this filtration step were considered genomes.

### 2.4 Taxonomic Assignment, Host Assignment, and Lifestyle Prediction

Data from Unified Human Gut Virome (UHGV; https://github.com/snayfach/UHGV) catalog was downloaded, and our predicted viral genomes were searched against it. BLASTn (Camacho et al., 2009) was used for identifying similar sequences by combining ≥ 95% identity with ≥ 500 bp length to calculate the coverage. If the query genome had an average ANI (BLASTn) of ≥ 95%, ≥ 70% and <70% with UHGV sequences, then the genomes were considered as identical, partially identical, and novel sequences, respectively (Chen et al., 2024). Metadata was fetched from UHGV for the identical and partially identical sequences (Table S1). PhaGCN (Shang et al., 2021) was used with default parameters for the taxonomic classification of the novel sequences. Furthermore, host assignment of the novel sequences was done by CHERRY (Shang & Sun, 2022). After this step, any sequence that was classified into any other realm than *Duplodnaviria* was excluded from further analysis.

BACPHLIP 0.9.6 (Hockenberry & Wilke, 2021) was used to predict the lifestyle of the phages. If the “Virulent” score was ≥ 0.9, then the phage sequence was marked as a Virulent phage. In the same way, any sequence with ≥ 0.9 “Temperate” score was considered a Temperate phage. Lifestyle of the remaining sequences was predicted with PhaTYP (Shang et al., 2023). Sequences that could not be predicted for virulent or temperate lifestyles, even with the latter tool, were marked as “Undetermined”.

### 2.5 Functional Annotation of Phage Sequences

The phage sequences were annotated in Pharokka (Bouras et al., 2023). Pharokka is a command-line-based pipeline that uses Prodigal (Hyatt et al., 2010) or Phanotate (Mcnair et al., 2019) for annotating bacteriophage sequences. The tool is inspired by Prokka (Seemann, 2014) and specializes in annotating bacteriophages. Here, Prodigal and the metagenomic mode (-p flag) were used for the annotation of phage sequences. The previously predicted high-quality genomes were used as the input for annotation.

### 2.6 Genome Clustering and Construction of Viral Proteomic Tree

The predicted proteins were clustered into phamilies (protein families) using phammseqs (Gauthier et al., 2022). These phamilies were then used to cluster the phage genomes according to their PEQ (proteomic equivalence quotient) index using phamclust (Gauthier & Hatfull, 2023). After that, the sequences in clusters were sorted depending on their genome completeness and then genome length. Sequences with the highest completeness and length were then selected from each cluster as representative sequences. A viral proteomic tree was built using these representative sequences with the prokaryotic viral sequences from Viral RefSeq (v230) (Pruitt, 2004) using ViPTreeGen (v1.1.3) (Nishimura et al., 2017). iTOL v7 was used for the annotation of the tree (Letunic & Bork, 2024).

### 2.7 Endolysin Prediction and Construction of Phylogenetic Tree

All proteins of any protein phamily that contained at least one annotated endolysin protein were marked as “Putative” endolysins. Then CD-HIT (-c 0.95 -n 5 -aS 0.8) was used to remove redundancy in the endolysin sequences. These non-redundant endolysin sequences were used in further analysis (Table S2). MAFFT v7.525 (Katoh, 2005) was used for the multiple alignment sequence, and the phylogenetic tree was built using FastTree 2.2 (Price et al., 2010) with default parameters. The tree was annotated with iTOL v7.

### 2.8 Domain Architecture Prediction

HMMER version v3.4 (Finn et al., 2011) was used to search the putative endolysins against the Pfam database for domain prediction. Hmmscan with default parameters was used in the search. While screening the sequences for better prediction, the threshold for all the e-values, including e-value, i e-value, and c e-value, was set to ≤0.0001. Again, predicted domains with the bit score and bias on the same order of magnitude were excluded. The threshold for bias was set to ≤10. Additionally, InterProScan (Hunter et al., 2009) and phmmer were used by sequence submission to validate the results from Hmmscan.

## 3. Results

### 3.1 Extraction of Phage Genomes from the Gut Metagenomes

A total of 9,141 human gut metagenome samples containing 169,736,757 contigs across 34 datasets from 20 different countries were analyzed. From these contigs, 592,534 viral sequences were predicted. After that, 21,690 phage genomes were extracted based on viral signatures and genome completeness. These genomes were checked for redundancy. Finally, 15,267 non-redundant high-quality and complete sequences were selected for further analysis.

The human gut metagenome samples were from 20 different countries. These countries are the US, Canada, Denmark, Spain, Germany, France, Russia, Estonia, Finland, India, Madagascar, China, Kyrgyzstan, Uganda, Cameroon, Italy, South Korea, Israel, Ireland, and the Netherlands. Among these countries, the US had the largest sample size (n = 4741; 51.87%) (Figure 1A).

**Figure 1:**
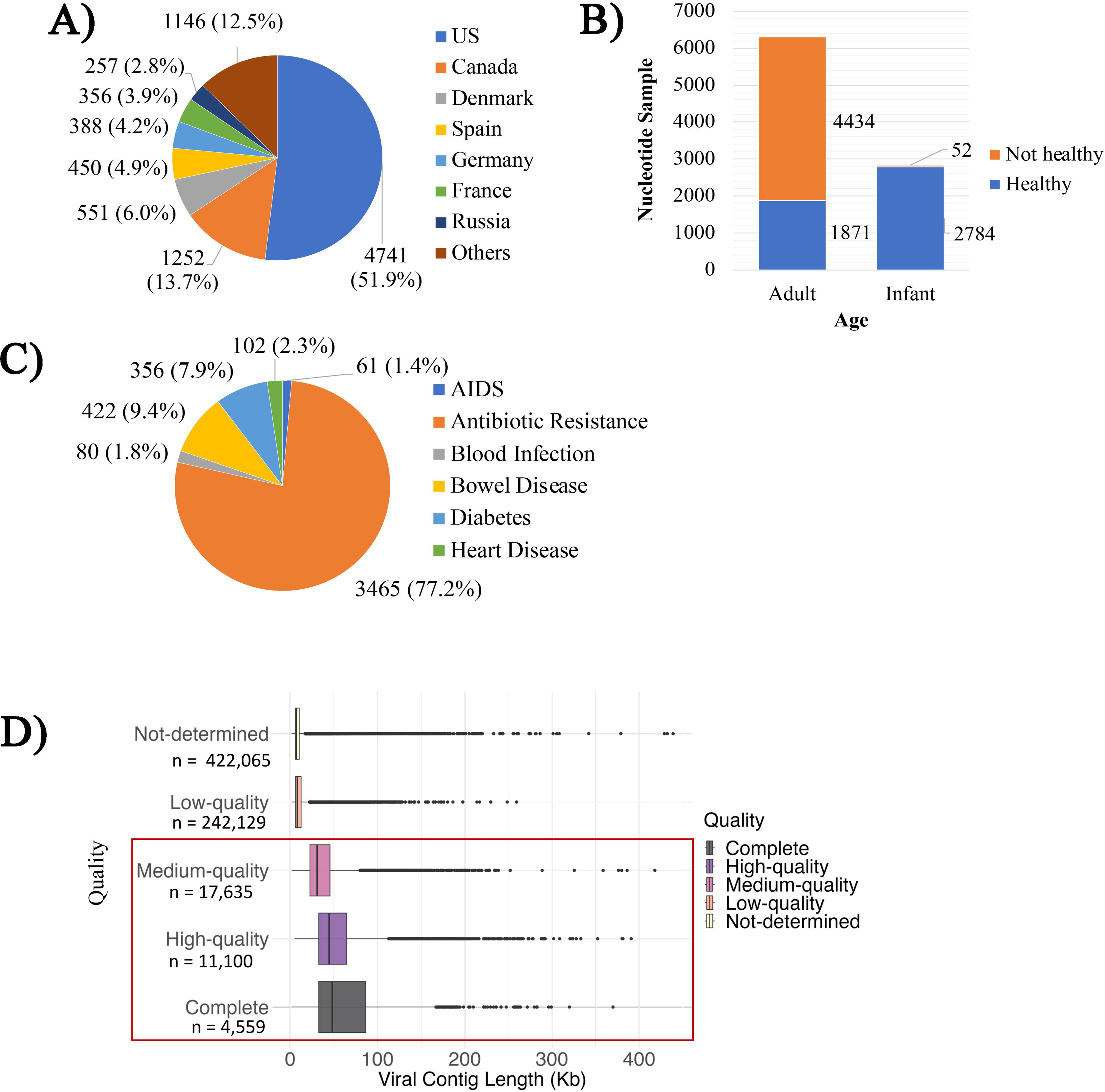
A) Global distribution of samples. More than half (51.87%) of the samples were collected from the United States alone. B) The human gut metagenomic sample distribution according to age and health. Blue and orange represent healthy and not healthy cohorts, respectively. The x-axis denotes the nucleotide sample, while the y-axis denotes the adult and infant cohorts, respectively. C) Sample distribution according to the disease in the previously mentioned not healthy cohort. D) chechV report summary of the gut phageome. The X-axis denotes viral contig length, and the Y-axis denotes the number of bacteriophage genomes.

The dataset included two age groups: adult (n = 6,305, 68.97%) and infant (n = 2836, 31.03%) (Figure 1B). Among the unhealthy population, antibiotic resistance samples were most prevalent (n = 3465; 77.24%) with most collected during FMT (Fecal Microbiota Transplant) treatment. Bowel diseases are the second largest category in the unhealthy population (n = 422; 9.41%), comprising Enterocolitis, Short Bowel Syndrome, Colitis, Inflammatory Bowel Disease, and Gastric Ulcer. The rest of the samples are from the population with heart disease (n = 102; 2.27%), blood infection (n = 80; 1.78%), Type-1 diabetes (n = 356; 7.94%), and AIDS (n = 61; 1.36%) (Figure 1C). The dataset also contained 17,635 medium-quality genomes (completeness 50–90%), 11,100 high-quality genomes (completeness >90%), and 4,559 complete genomes (completeness 100%) (Figure 1D).

### 3.2 Prediction of Phage Lifestyle, Their Gene Count, GC Content, and Genome Size Distribution

A total of 15,267 genomes were identified, comprising 12,702 linear (83.20%) and 2,565 circular (16.80%) sequences. Bacteriophage sequences were analyzed for lifestyle prediction, identifying 7,667 virulent and 6,691 temperate phages, while 138 genomes (0.95%) could not be classified (Figure 2A). Genome size distribution showed a bimodal pattern with a long tail, with most genomes ranging from ∼25 kbp to ∼75 kbp, averaging at ∼48 kbp (Figure 2B). Genome size exhibited a strong positive correlation with gene count (Pearson r = 0.9322, P < 0.001), indicating that larger phage genomes encode more genes, with an average of ∼157 genes per genome. Conversely, genome size showed a moderate negative correlation with GC content (Pearson r = –0.3020, P < 0.001), suggesting that GC content tends to decrease with increasing genome size, although the overall GC content remained relatively stable at ∼43% across phages. A weak negative correlation was also observed between genome size and coding efficiency (Pearson r = –0.2049, P < 0.001), indicating a slight decline in coding efficiency as genome size increases (Figure 3).

**Figure 2:**
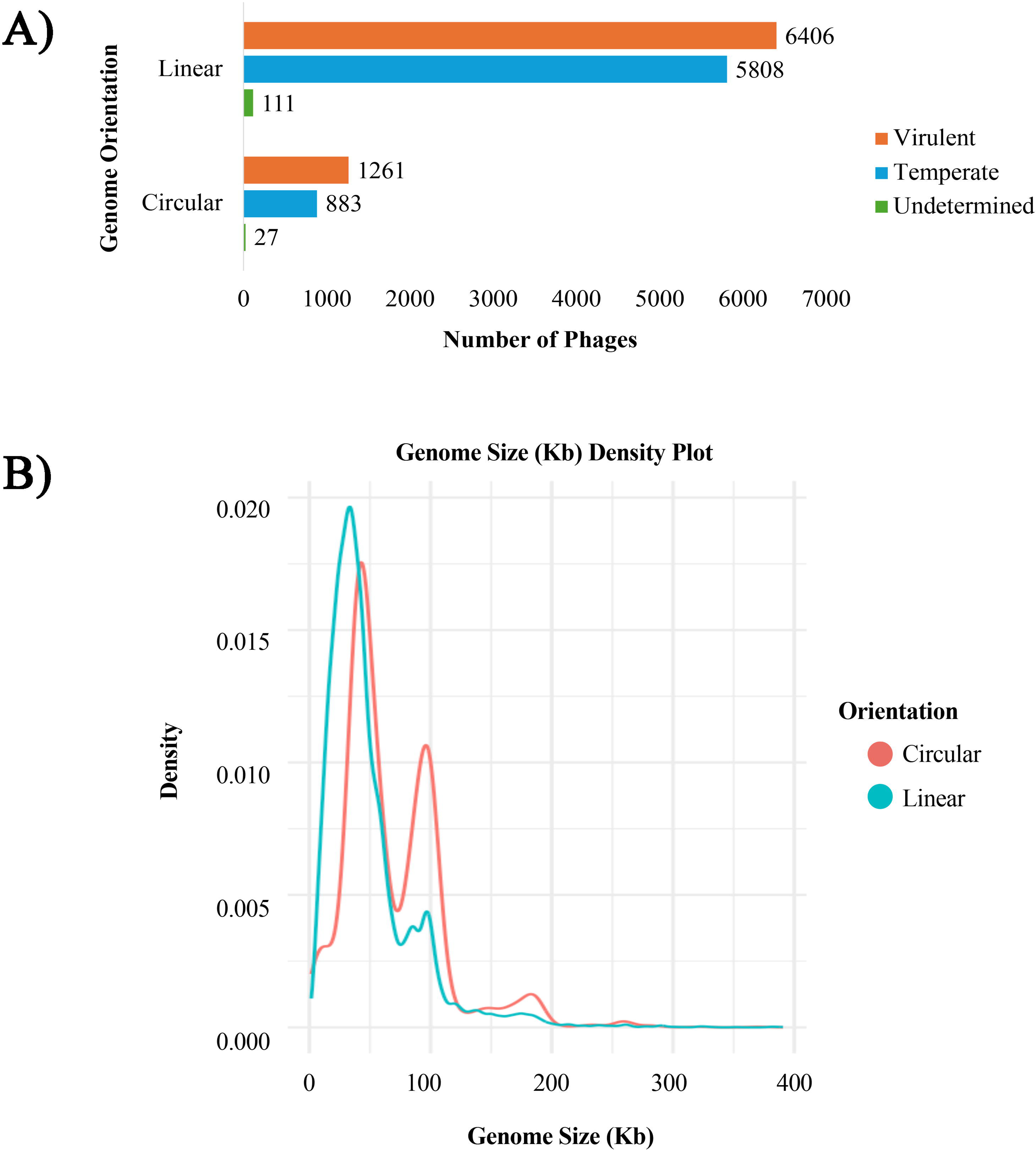
A) A bar plot showing the number of phages maintaining a temperate and virulent lifestyle. The x-axis denotes the lifestyle, while the y-axis denotes the number of phages. Orange, blue, and green colors correspond to circular, linear, and undetermined genome orientation, respectively. B) The density plot of phage genome size in the phageome showing a bimodal distribution with a long tail. Most of the detected bacteriophages were between the range of ∼25 kbp to ∼75 kbp. Again, sequences with more than 250 kbp are phages with large heads and large genomes.

**Figure 3:**
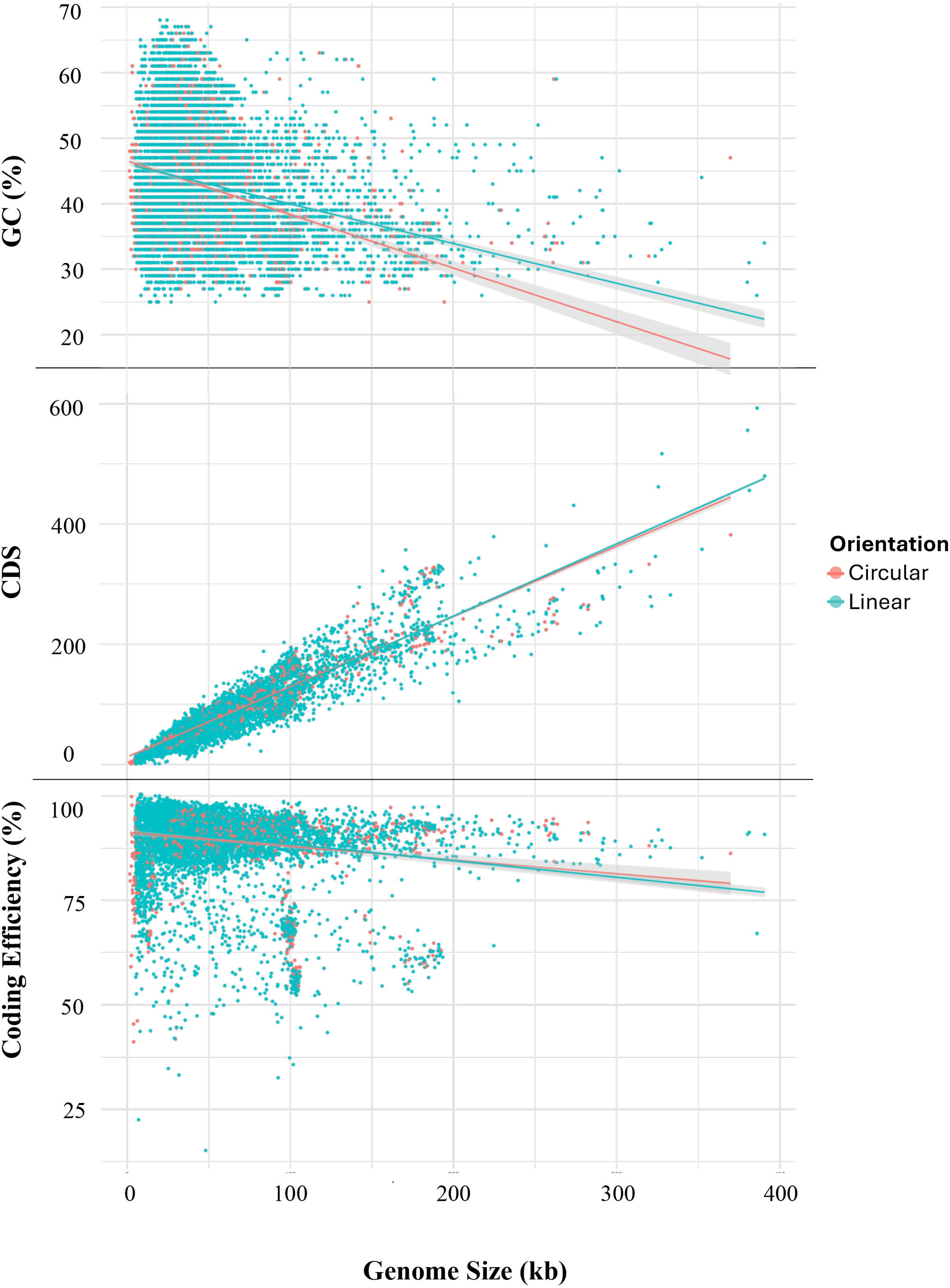
A scatterplot showing the correlations of GC content, CDS, and Coding efficiency with Genome size. The correlation between GC content and genome sizes was moderately negative (Pearson r = - 0.3020, P < 0.001). Despite the negative correlation, on average, ∼43% GC content per genome was observed, with a low amount of variation. Between the genome sizes and the number of genes in the phages, a highly positive statistically significant correlation was observed (Pearson r = 0.9322, P < 0.001), averaging at ∼157 genes per genome. On the other hand, there was a weak correlation between genome size and coding efficiency (Pearson r = - 0.2049, P < 0.001).

### 3.3 Lifestyle Prediction, Taxonomic Assignment, and Abundance Analysis

Comparison with the UHGV database revealed 4,825 identical, 3,084 partially identical, and 7,323 novel sequences. Taxonomic analysis assigned 2,243 phages to 24 families, with *Crassviridae* being the most abundant (n = 941; 41.95%), further classified into *Alpha/Gamma-crassviridae* (n = 321), *Beta-crassviridae* (n = 90), *Delta-crassviridae* (n = 329), *Epsilon-crassviridae* (n = 13), and *Zeta-crassviridae* (n = 188). Next came *Flandersviridae* (n = 359, 16.01%), *Suoliviridae* (n = 189, 8.43%), *Intestiviridae* (n = 126, 5.62%), *Autographiviridae* (n = 93, 4.15%), *Steigviridae* (n = 92, 4.10%), *Gratiaviridae* (n = 91, 4.06%), *Crevaviridae* (n = 64, 2.85%), *Salasmaviridae* (n = 59, 2.63%), *Quimbyviridae* (n = 43, 1.92%), *Orlajensenviridae* (n = 39, 1.74%), and several less represented families grouped as others (n = 147, 6.55%). The “Others” category included are *Aliceevansviridae* (n = 29, 1.29%), *Drexlerviridae* (n = 24), *Peduoviridae* (n = 17), *Duneviridae* (n = 14), *Schitoviridae* (n =12), *Guelinviridae* (n = 12), *Winoviridae* (n = 12), *Straboviridae* (n = 10), *Rountreeviridae* (n = 9), *Casjensviridae* (n = 6), *Herelleviridae* (n = 1), and *Zobellviridae* (n = 1). Notably, the largest portion of sequences, mostly circular and virulent, remained unannotated (n = 12,253; 84.53%) and were grouped as UC_*Caudoviricetes* (Unclassified_*Caudoviricetes*). Additionally, three jumbo phages were identified (Figure 4A). All phage sequences were clustered into 825 clusters and 736 singletons and their representative sequences, along with singletons, formed clusters with prokaryotic viral reference sequences in the viral proteomic tree (Figure 4B).

**Figure 4:**
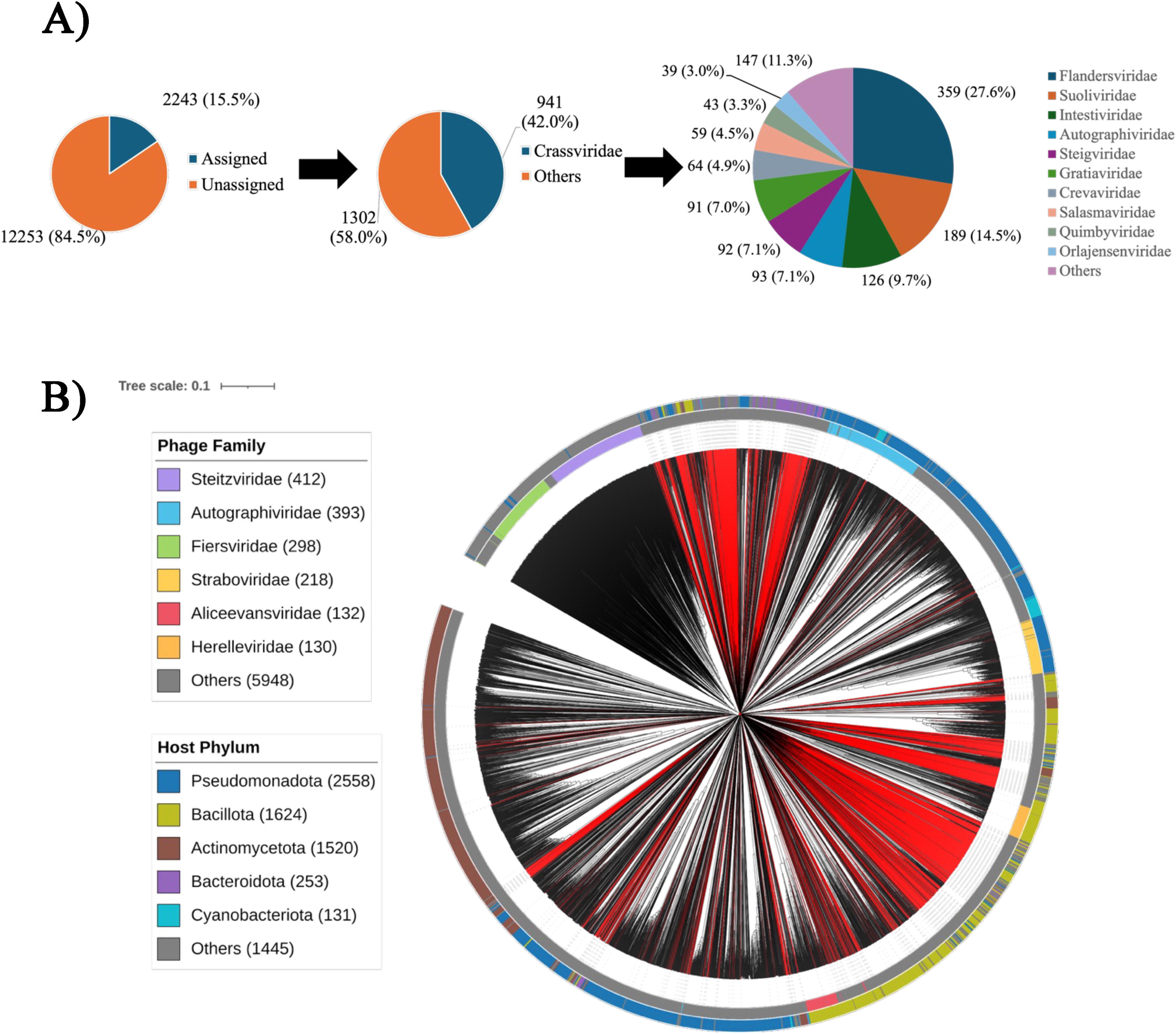
A) A pie chart showing the abundance of globally distributed bacteriophages in the human gut microbiome. More than four-fifths (84.53%) of the genomes were not classified. B) A viral proteomic tree of bacteriophage genomes. The inner circle denotes the phage family, and the outer circle indicates the host phylum. Again, the red branches represent the sequences of this study, and the black branches show the sequences of RefSeq viral sequences.

### 3.4 Host-Phage Association Analysis

Host–phage association analysis assigned phage hosts at the species level, identifying 19 unique phyla, 368 genera, and 1,140 species. Over two-fifths of phages (n = 6,157; 41.06%) were associated with hosts from the *Bacillota* phylum, followed by *Bacteroidota* (n = 3,511; 23.41%), *Pseudomonadota* (n = 1,099; 7.33%), and *Actinomycetota* (n = 985; 6.57%). In total, 12,485 phage genomes (83.26%) were successfully assigned to hosts, while 2,511 remained unassigned (Figure 5A). All phage families except *Zobellviridae* had at least one member predicted to infect Bacillota, with *Crassviridae* showing the highest abundance within this group (n = 345). Most *UC_Caudoviricetes* (n = 5352) were linked to *Bacillota*. Comparable diversity was observed in *Bacteroidota*, which served as hosts for 23 phage families, the same number as *Bacillota*, while *Pseudomonadota* and *Actinomycetota* hosted phages from 19 and 18 families, respectively (Figure 5B). The overall phage–host associations were further illustrated using a Sankey diagram (Figure 5C).

**Figure 5:**
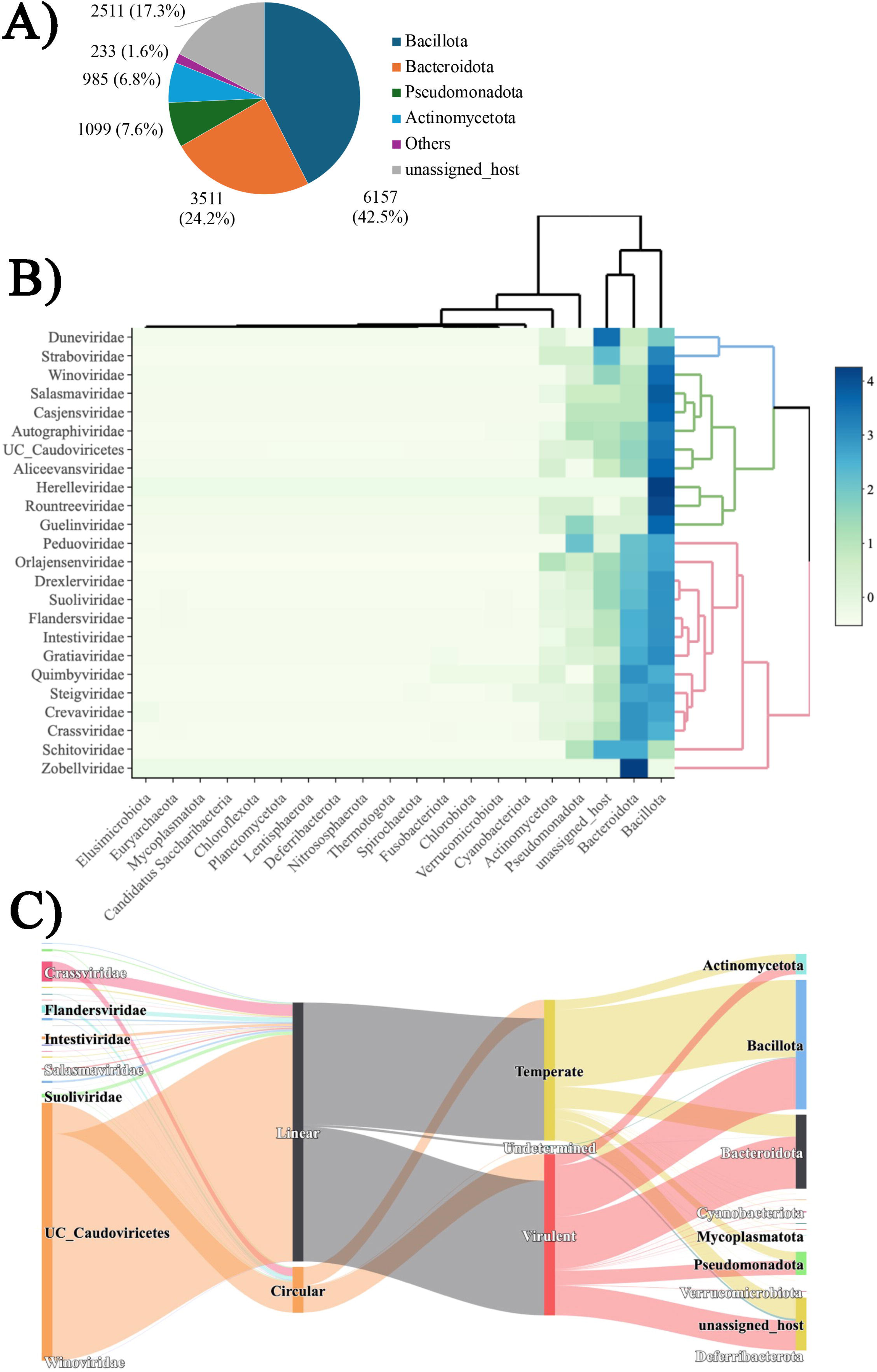
A) A pie chart showing the distribution of hosts predicted for the bacteriophages in the human gut microbiome. Almost half (42.5%) of the predicted hosts were *Bacillota*. B) A heatmap showing the bacteriophage-host bacteria association in the human gut microbiome. The four major phyla of bacteria (*Bacillota, Actinomycetota, Pseudomonadota, Bacteroidota)* in the human gut microbiome showed the most hits for predicted hosts. C) A Sankey graph showing the bacteriophage-host bacteria association according to their lifestyle and genome orientation.

### 3.5 Protein Diversity of Gut Phageome

The phageome encoded a total of 1,012,676 proteins, of which nearly four-fifths (n = 799,405; 78.94%) remained unannotated, while the remaining proteins were assigned to various functional classes. Among the annotated proteins, those involved in DNA, RNA, and nucleotide metabolism were the most abundant (n = 70,972; 7.01%). Proteins related to phage–host interactions were relatively limited in number; for example, morons, auxiliary metabolic genes, and host takeover proteins totaled 6,883, while only 5,546 proteins were annotated with integration or excision functions (Figure 6A). The analysis identified 25 VFDB hits, primarily from *Salmonella* spp. and *Shigella* spp., with additional hits from *Streptococcus* spp. A total of 19 antibiotic resistance genes were detected, 9 of which corresponded to *Enterococcus faecium.* Overall, the proteome clustered into 140,668 protein families (phamilies).

**Figure 6:**
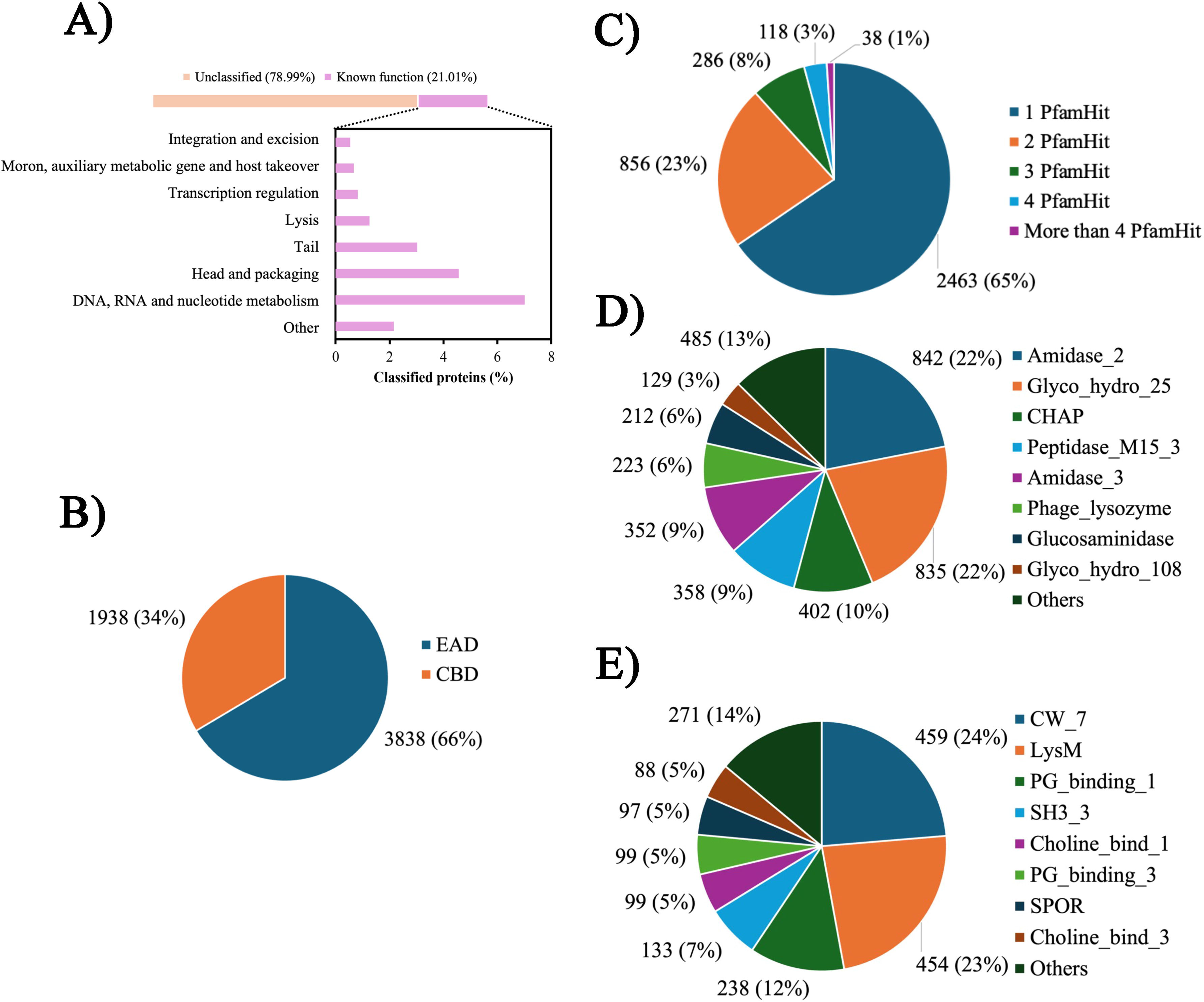
Distributions of EAD & CBD, Pfam hits, and all predicted endolysin domains. A) Annotation of proteins of gut phageome. Almost two-fifths of the proteins could not be classified. B) A pie chart showing the percentage of Enzymatically Active Domains (EAD) and Cell Binding Domains (CBD) corresponding to the blue and orange colors, respectively. C) A pie chart showing the number of PfamHits for each putative endolysin sequence. D) The percentage of all detected endolysin EADs. E) The percentage of all detected CBDs.

### 3.6 Domain Architectures, and Phylogeny of Endolysins

Through detailed curation, a refined dataset of 3,794 phage lytic proteins (Table S3) was generated, comprising 296 phamilies and 266 Orphams (singletons). The distribution of enzymatically active domains (EADs) and cell-binding domains (CBDs) was 66% and 34%, respectively (Figure 6B; Figure A1). Domain annotation revealed that 2,463 proteins contained a single significant Pfam hit, while 856 had two hits. Additionally, 286, 118, 26, 5, and 6 proteins contained three, four, five, six, and seven Pfam hits, respectively. Remarkably, one protein contained 15 domains (Figure 6C). The most frequently occurring EAD domains among endolysins were Amidase_2 (n = 842) and Glyco_hydro_25 (n = 835) (Figure 6D) and CW_7 (n = 459, 24%) and LysM (n = 454, 23%) were the most abundant CBDs (Figure 6E). Single-domain proteins primarily contained Amidase_2 (n = 561; 22.75%), Glyco_hydro_25 (n = 426; 17.27%), and Peptidase_M15_3 (n = 307; 12.45%) (Figure 7A).

**Figure 7:**
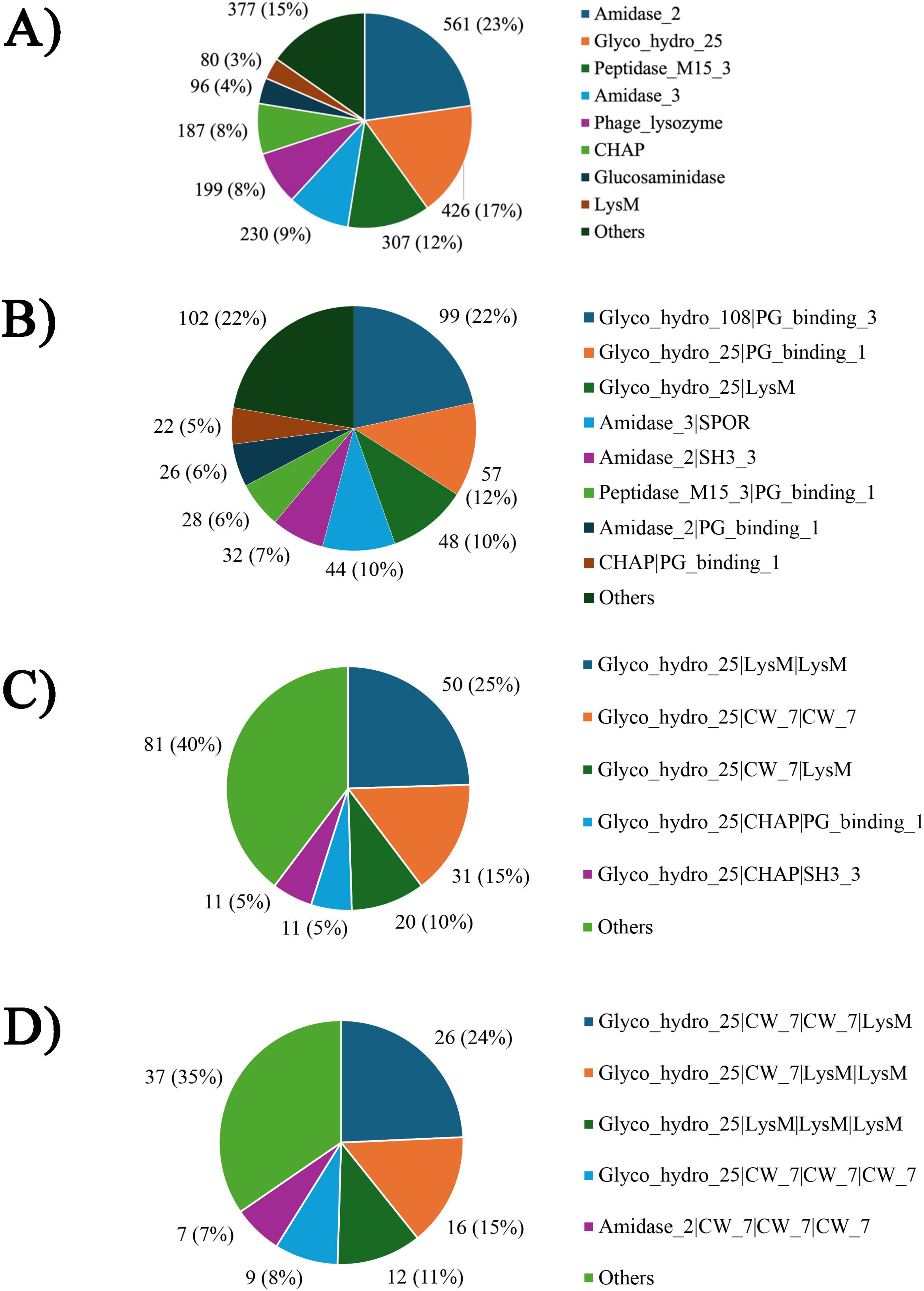
Distributions of predicted domains in Endolysins. A) The percentage of detected endolysins with single domains. Amidase_2 occupied more than one-fifths (23%) abundance among the other single-domain catalytic endolysins. B) The percentage of detected endolysins with two domains. Glyco_hydro_108 and PG_binding_3 endolysin were the most abundant (22%). C) The percentage of detected endolysins with three domains. Glyco_hydro_25, LysM, and LysM, a monocatalytic endolysin, were the most abundant (25%). D) Endolysins with four domains. Glyco_hydro_25, CW_7, CW_7, and LysM were the most abundant (24%).

Proteins with two Pfam hits displayed diverse EAD–CBD combinations. The most common architecture consisted of Glyco_hydro_108 (EAD) and PG_binding_3 (CBD) (n = 99; 22%). Other prevalent combinations included Glyco_hydro_25 (EAD) with PG_binding_1 (CBD) (n = 57; 12%) and Glyco_hydro_25 (EAD) with LysM (CBD) (n = 48; 10%). Additional notable patterns included Amidase_3 with SPOR, Amidase_2 with SH3_3, and Peptidase_M15_3 with PG_binding_1 (Figure 7B). Proteins with three domain hits exhibited either bicatalytic or monocatalytic multimodular arrangements. Among bicatalytic architectures, Glyco_hydro_25–CHAP–PG_binding_1/SH3_3 (n = 11; 5%), CHAP–Amidase_2–SH3_5 (n = 5; 2%), and CHAP–NAGPA–SH3_3 (n = 4; 2%) were common. For monocatalytic proteins, the most frequent architecture was Glyco_hydro_25–LysM–LysM (n = 50; 25%). Other major patterns included Glyco_hydro_25–CW_7–CW_7 (n = 31; 15%) and Glyco_hydro_25–CW_7–LysM (n = 20; 10%) (Figure 7C).

Endolysins with four domains displayed monocatalytic, bicatalytic, and multicatalytic architectures. Monocatalytic forms commonly featured Glyco_hydro_25 EADs with CBDs such as CW_7 or LysM, and Amidase_2 or Amidase_5 EADs paired with CW_7, SPOR, Choline_bind_1, or Choline_bind_3. Bicatalytic four-domain proteins included Phage_lysozyme2–CHAP–LysM–LysM and Glyco_hydro_25–LGFP–LGFP–NLPC_P60 (n = 2; 2%). Multicatalytic proteins included combinations such as Peptidase_C39_2–Peptidase_C39_2–Phage_lysozyme2–NLPC_P60 (n = 2; 2%), and Phage_lysozyme2–Amidase_5–Glucosaminidase–CHAP (n = 2; 2%). A unique protein consisted of four LysM domains. Overall, a modular organization was evident across these endolysins (Figure 7D).

Proteins containing five domains were mainly monocatalytic (n = 15; 56%), featuring EADs such as Glyco_hydro_25, Amidase_2, Glucosaminidase, or Amidase_5 combined with CBDs like Choline_bind_1, Choline_bind_3, LysM, SH3_3, and CW_7. Bicatalytic proteins (n = 11; 41%) combined catalytic domains such as Glucosaminidase, Glyco_hydro_25, Phage_lysozyme2, or CHAP with various CBDs. Only one protein exhibited EADs flanking the CBDs (Glyco_hydro_25–LGFP–LGFP–NLPC_P60; n = 1). A single multicatalytic five-domain protein contained CHAP–Phage_lysozyme2–Amidase_2–SPOR–CW_7.

Six-domain endolysins were rare (n = 5) and mostly monocatalytic, containing EADs such as Amidase_2, Amidase_3, or CHAP, paired with Choline_bind_1, Choline_bind_3, Choline_bind_4, or ChW CBDs. One bicatalytic protein carried Glucosaminidase, CHAP, and four CW_7 domains. Additionally, five seven-domain proteins—all monocatalytic—were identified, with EADs such as Amidase_3 or Amidase_2, combined with CBDs like ChW or Choline_bind_4. The most complex protein identified contained 15 Pfam domains, including Glyco_hydro_25, thirteen Choline_bind_4 domains, and YkuD. This bicatalytic protein had EADs positioned at both termini and CBDs clustered centrally.

Phylogenetic reconstruction showed that endolysins belonging to the same phamilies predominantly clustered together, highlighting shared evolutionary origins and conserved structural modularity across groups (Figure 8).

**Figure 8:**
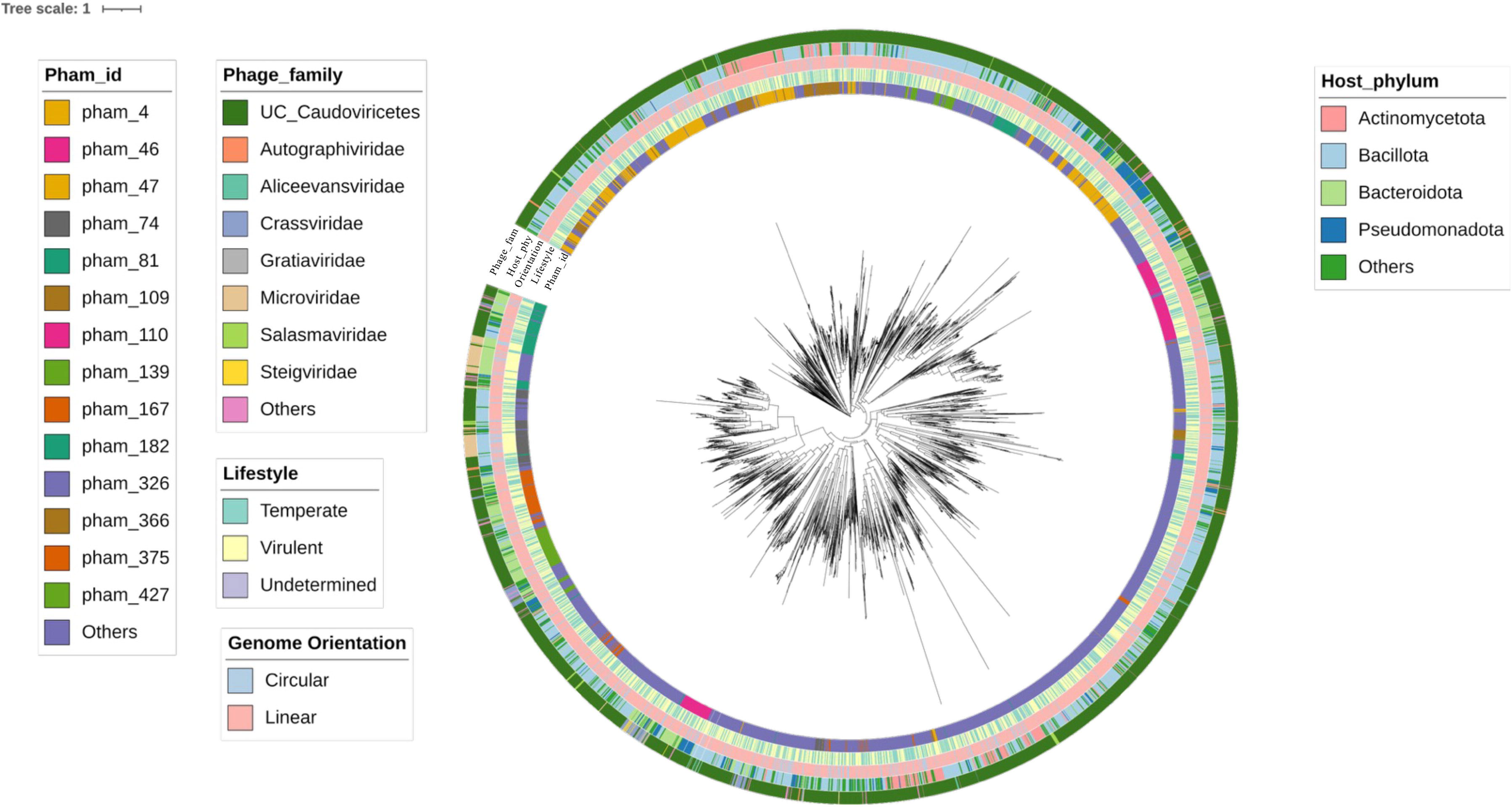
Radial maximum-likelihood phylogenetic tree of endolysin proteins of human gut phageome. Concentric rings (inner to outer) denote phamily, phage family, phage lifestyle, phage genome orientation, and host bacterial phylum. Branch lengths represent substitutions per site. The figure illustrates the broad evolutionary and ecological diversity of gut phage-encoded endolysins.

## 4. Discussion

Phages constitute the majority of the viral fraction within the gut microbiome, reaching abundances greater than 10¹L gL¹ (Hoyles et al., 2014; Shkoporov et al., 2018). Their strong ecological influence shapes both the composition and function of the human gut microbial community in health and disease (Townsend et al., 2021). As antibiotic resistance within gut bacteria becomes increasingly widespread, traditional treatment strategies face growing limitations. This has accelerated interest in precision antimicrobials such as phageLencoded endolysins, which exhibit potent, species-level specificity (Gontijo et al., 2021; Mondal et al., 2020; Rahman et al., 2021). Using extensive gut metagenomic datasets, this study characterizes the gut phageome with a particular focus on endolysins and their structural and functional diversity. Our findings uncover numerous previously uncharacterized enzymes and expand the resources available for developing targeted, next-generation antimicrobials for gastrointestinal pathogens, including antibiotic-resistant strains.

Our analysis identified, 15,267 high-quality, non-redundant genomes after stringent filtering. Genome size displayed a bimodal distribution, dominated by genomes between 25 and 75 kbp but also including a long tail of larger genomes with three jumbo phages. This distribution mirrors earlier global virome studies (Hatfull, 2008; Paez-Espino et al., 2016), while also highlighting the persistence of rare large phages in human populations. Genome size correlated strongly with gene count, as also reported previously (Gao et al., 2020; Ha & Denver, 2018), reinforcing that genome expansion reflects increased coding capacity rather than accumulation of nonfunctional DNA. Larger genomes tended to have lower GC content and slightly reduced coding density, consistent with patterns linked to phage–host adaptation (Almpanis et al., 2018). Together, these trends reveal a two-tier phage community: abundant, compact phages optimized for efficient replication, and more complex, gene-rich phages positioned for broader functional capabilities.

Taxonomic analysis assigned the bacteriophages into 24 families according to the current ICTV classification (Turner et al., 2021, 2023). *Crassviridae* was found to be the most abundant phage family in the gut, aligning with previous studies (Dutilh et al., 2014; Guerin et al., 2018). However, nearly 85% of phages remained unassigned, and more than 7,000 sequences appeared novel. Moreover, proteomic tree analysis pointed out that assigned phages often failed to form monophyletic clades, underscoring previously unseen diversity. These findings emphasize the need for deeper investigation into gut phageome structure, classification, and evolution.

Host prediction succeeded for more than 83% of genomes and revealed a striking ecological pattern: 41% of all phages targeted *Bacillota*, followed by *Bacteroidota*, *Pseudomonadota*, and *Actinomycetota*. This *Bacillota* dominance is consistent with earlier work (Nayfach et al., 2021) and reflects the phage pressure exerted on both beneficial and pathogenic members of this phylum. Given that *Bacillota* include some of the gut’s most important commensals, such as *Faecalibacterium* and *Roseburia*, as well as problematic pathogens like *Clostridioides difficile* and vancomycin-resistant *Enterococcus faecium* (Chisti et al., 2021). The prevalence of gut phages that specifically target *Bacillota* provides a framework for developing precision phage-based therapies. For instance, phages and their associated endolysins may be engineered to selectively eliminate pathogens like *C. difficile* while preserving beneficial commensals (Mondal et al., 2020).

Our dataset revealed a nearly equal representation of virulent (lytic) and temperate (lysogenic) phage lifestyles. This balance persisted across both the more prevalent linear-genome phages (∼80%) and the less common circular-genome phages. Furthermore, specific virulent lineages (e.g., crAss-like phages) maintain substantial populations (Shkoporov et al., 2019; Shkoporov & Hill, 2019). This equilibrium highlights a shared co-evolutionary dynamic across viral genomes. Virulent phages impose strong top-down regulation, driving rapid bacterial turnover and influencing community composition. In contrast, temperate phages, which integrate into host genomes, function as long-term genetic reservoirs that facilitate horizontal gene transfer and enable bacterial evolution (Howard-Varona et al., 2017).

The presence of both lifestyles in linear and circular genomes, such as lambda phages and families like *Microviridae* and *Inoviridae*, suggests that the balance between lytic and lysogenic strategies is a fundamental ecological feature of gut phages. This duality helps maintain microbial stability while enabling continuous evolutionary adaptation (Casjens & Gilcrease, 2009; Roux et al., 2012).

The most remarkable outcome of this study is the creation of a comprehensive catalog of 3,794 gut phage endolysins grouped into nearly 300 families. Beyond the expected single-domain enzymes (Criel et al., 2021), the dataset reveals extensive structural complexity, including dozens of lysins with four or more domains. Several proteins carried up to seven domains, and an exceptional 15-domain lysin was identified, containing catalytic domains flanking 13 consecutive binding modules. Such extreme modularity is unprecedented, far exceeding previously reported natural lysins (Fernández-Ruiz et al., 2018) and even surpassing the level of complexity typically achieved through synthetic engineering (Briers et al., 2014; Pottie et al., 2024; Walsh et al., 2021). Domain combinations frequently involved well-known catalytic modules such as Amidase_2 and Glyco_hydro_25 linked with binding repeats like LysM, CW_7, and Choline_bind. These findings highlight the gut as an evolutionary hotspot for creating highly modular lytic enzymes, offering a vast natural reservoir for therapeutic design. As shown in recent work (Wang et al., 2024), even modest expansions in modular architecture can improve stability and broaden activity. The discovery of such extreme domain multiplicity indicates that phages have explored functional landscapes well beyond current engineering capabilities.

Together, the results relating to genome architecture, host distribution, and endolysin modularity point toward a coherent evolutionary logic. Larger phage genomes encode a greater number of genes, including the blueprints for complex lytic enzymes. These phages are overwhelmingly linked to Gram-positive *Bacillota* hosts, whose multilayered, resilient peptidoglycan walls pose significant lysis challenges. In direct response to this ecological pressure, endolysins have evolved increasingly modular structures, integrating multiple catalytic and binding domains to enhance lytic efficiency. This interplay among genomic potential, host ecology, and enzyme architecture provides a logical foundation for therapeutic innovation, shifting the focus from *de novo* design to mining the superior, naturally optimized resources available in the gut phageome.

We acknowledge certain inherent limitations in computational metagenomic analysis. Assembled metagenomes can occasionally produce chimeric proteins, potentially inflating domain counts, and reference databases inherently limit the accuracy of domain annotation. While computational predictions of hosts and lifestyles are getting increasingly accurate, they ultimately require culture-based confirmation. Furthermore, our dataset reflects a bias toward cohorts exposed to antibiotics and those treated with fecal microbiota transplantation (FMT), meaning the full viral diversity of a pristine, healthy gut population may not have been fully captured. Future work will be critical for translating these findings into clinical tools. Structural studies are essential to elucidate the cooperative mechanisms of multi-domain lysins. Subsequently, synthetic pipelines should be established to recombine prevalent modules (e.g., Amidase_2, Glyco_hydro_25 with LysM or CW_7) into optimized therapeutic endolysins. Finally, expanding sampling into underrepresented global populations will be necessary to determine if this extreme modularity is a gut-specific phenomenon or a general strategy of phage evolution.

## 5. Conclusion

This study provides the most extensive genomic and functional characterization of the human gut phageome to date. We demonstrate a diverse array of bacteriophages with a clear dominance toward infecting *Bacillota* and *Bacteroidota*, illustrating the complexity of the gut virome ecology. Crucially, the identification of approximately 3,800 highly modular endolysins, including structures of unprecedented complexity, establishes the gut phageome as a prolific source of novel natural antibacterial agents. The existence of both basic single-domain lysins and intricate multi-domain structures illustrates nature’s sophisticated mechanisms for cell wall degradation. These findings significantly enhance our fundamental understanding of gastrointestinal phage ecology and provide a critical foundation for the development of next-generation, phage-based therapeutics targeting antibiotic-resistant bacteria.

## Acknowledgement

Not applicable

## Ethics approval

None required

## Data Availability Statement

The raw data supporting the findings of this study are available from the corresponding author upon reasonable request.

## Supplementary Figure Legends

**Supplementary Figure 1**

Total distribution of endolysin domains in the human gut phageome.

