## Supplementary figures and images for "Human Gut Phageome Analysis Uncovers Thousands of Highly Modular Endolysins"

### Supplemental Figure 1

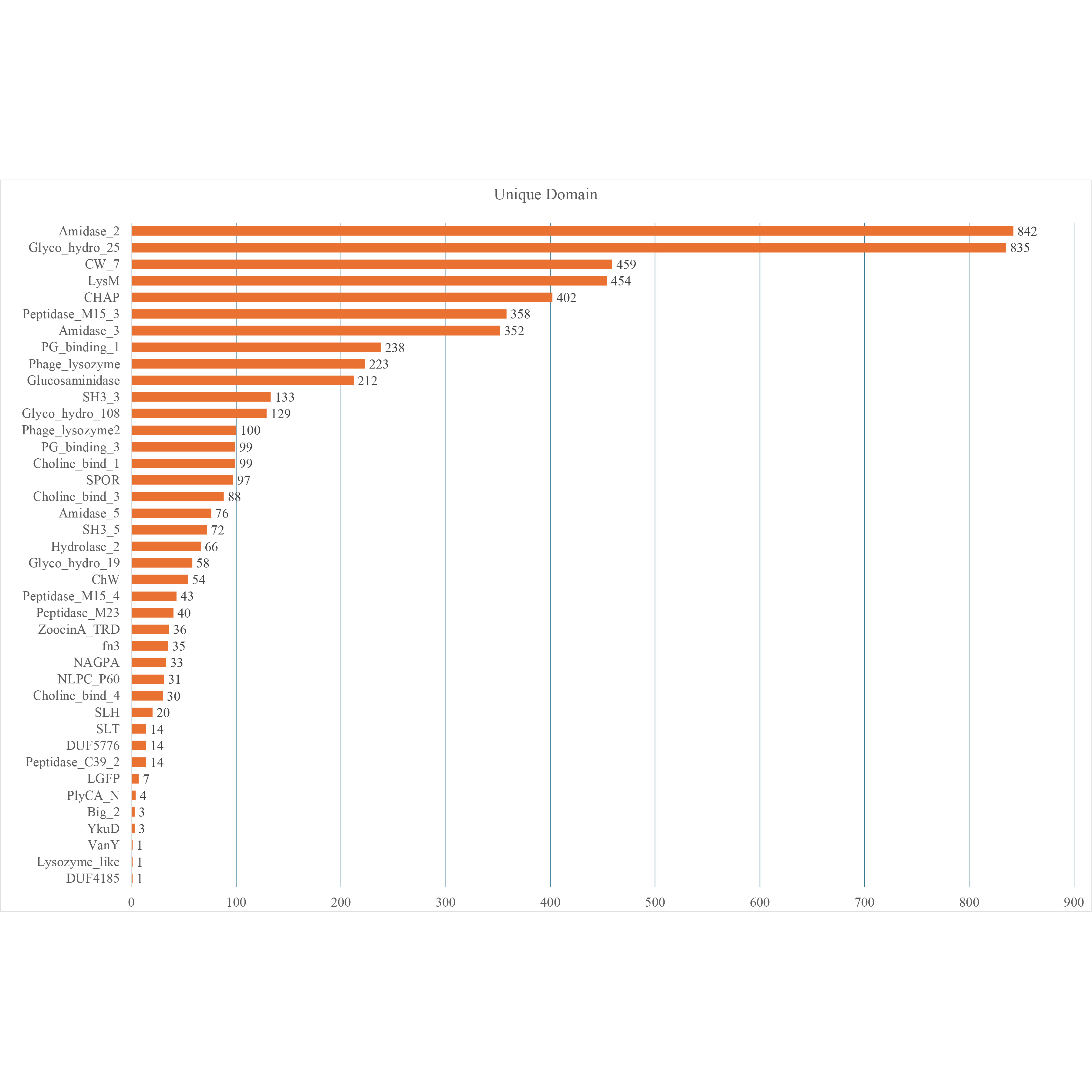
